# Discovery and characterisation of OMVs produced by the bee gut microbiota

**DOI:** 10.64898/2026.04.19.719495

**Authors:** Rodney P. Eyles, Waldan K. Kwong

## Abstract

Bacteria use diverse mechanisms to interact with each other and with eukaryotic hosts, thereby shaping microbiome composition and influencing host health. One of these mechanisms is the production of outer membrane vesicles (OMVs), nanoscale structures that bud off from bacterial cells into the extracellular space. OMVs can deliver bioactive cargoes, including enzymes, RNA and DNA, enabling functions such as cell-to-cell communication, nutrient acquisition and immunomodulation. However, the role of OMVs in beneficial host-associated microbiomes remains unclear. Here, we investigated OMV production in the gut bacteria of the western honey bee (*Apis mellifera*), which forms a highly conserved and stable microbial community. Using electron microscopy, fluorescence labelling, and nanoparticle tracking analysis, we detected OMV production in every gram-negative species of the bee gut microbiota that we investigated. Vesicles were observed in gut contents of wild and laboratory-inoculated bees, but absent in bees lacking a microbiota. OMVs contained nucleic acids, with more RNA than DNA. Bacterial strains varied in OMV properties, including abundance, size, and zeta potential. These findings indicate that OMVs are likely significant mediators of interbacterial and host-microbe interactions in the bee gut.

## 1. Introduction

Gut microbiomes are largely comprised of a core set of resident bacteria which form stable, structured communities that are highly adapted to the physical and chemical environment of the host. Gut microbes form symbiotic relationships with their host and contribute to its wellbeing, by increasing nutrient acquisition via breakdown of otherwise indigestible food material, strengthening and protecting the host gut lining, and boosting colonisation resistance from foreign or pathogenic bacteria [1]. Conversely, disruption to the gut microbiome (dysbiosis), is associated with a broad range of negative host health effects beyond those associated with digestion [2–4]. To maintain gut microbiome homeostasis, bacteria use a variety of signalling and secretion mechanisms to interact with each other as well as their hosts. One of the least well-understood modes of interaction are through outer membrane vesicles (OMVs). These are nanoscale extracellular structures derived through budding of the bacterial outer membrane. They can perform a variety of roles including iron acquisition [5], delivery of toxins [6], facilitation of horizontal gene transfer [7] and as antimicrobial agents through, for example, acting as decoys against phage predation [8, 9]. OMVs can deliver various functional cargoes such as DNA, RNA and proteins, including toxic proteases and lipases [2, 6, 10–12]. They may also directly affect gene expression within target cells through transfer of noncoding RNAs [13]. Research to date has focused on OMVs derived from pathogenic bacteria and their impact on the host, while less attention has been placed upon their role in mutualistic relationships or in maintaining homeostasis among the resident gut microbiota. Developing a detailed understanding of the signalling factors contributing to microbiome homeostasis is challenging as the composition of most gut microbiomes is complex and may contain thousands of bacterial species [14–16]. In contrast, the gut microbiome of the western honey bee, *Apis mellifera*, consists of a small number (8–10) of vertically transmitted [17] bacterial genera, making up ~95% of its total population [18]. This composition is largely stable and resistant to displacement by non-core species [19]. This relative simplicity makes it an ideal model for gut microbiome research.

The majority of the bee gut microbiota reside in the distal compartments of the gut. The pylorus and ileum sections are dominated by gram-negative bacteria, including *Frischella perrara, Gilliamella apicola, Gilliamella apis*, and *Snodgrassella alvi*; in contrast, the rectum is predominantly colonised by the gram-positive *Bifidobacterium* spp., *Bombilactobacillus* spp., and *Lactobacillus* spp. Twelve distinct bacterial secretion systems, including OMVs, have been classified to date [20] and several of these have been bioinformatically predicted to exist in the genomes of the *A. mellifera* gut microbiome. Only type 5 and type 6 secretion systems have been functionally characterized, in *S. alvi* [20, 21] and *F. perrara* [22]. Given the ubiquitous nature of gram-negative associated OMVs and their multifaceted functions, detection and characterisation of OMVs in the bee gut is an important step to revealing novel mechanisms contributing to microbiome colonisation and homeostasis. In this study we show that OMVs are abundantly produced by each gram-negative member of the resident bee microbiota, and that they are produced *in vivo*. We also establish baseline quantity, size and zeta potential for OMVs from multiple species, under differing growth conditions and show that they can carry nucleic acid cargo.

## 2. Materials and Methods

### (a) Bacterial culture and OMV collection

We examined members of the core bee gut microbiota for the production of OMVs. A complete list of strains and culture conditions used in this study is found in Table S1. For *in vitro* assays, liquid cultures were filtered through 0.2 μm pore membranes and centrifuged at 110,000–120,000 × *g*. The pellet was resuspended in dH_2_O and spun in a 50 kDa molecular weight cut-off filter at 3,000 × *g* to remove particles <50 kDa. Retentate containing the OMVs were collected and stored at 4°C. For *in vivo* assays, ilea contents from multiple adult worker bees colonised with *S. alvi* wkB2 or taken directly from the hive were collected and pooled, resuspended in dH_2_O and then treated as per the *in vitro* assays. Detailed methods are provided in Supplementary Methods.

### (b) Electron microscopy

Gut ileum tissue sections, bacterial cells, and purified OMVs (from bacterial cultures and gut ilea contents) were visualised with transmission electron microscopy (TEM) using negative staining. Three-dimensional (3D) tomography imaging was done using a tilt series collected at 23,000× magnification at 1° increments between −55° and 55°. Unstained, vitrified OMVs were used for cryo-TEM imaging. Samples were prepared using standard protocols as detailed in Supplementary Methods.

### (c) Confocal imaging of OMV-associated nucleic acids

OMVs from *G. apis* ESL0172 were treated with DNase I and RNase A to remove non-luminal nucleic acids. DNA or RNA spike-ins were used to determine digestion effectiveness. For OMV-DNA co-localisation, OMV samples were stained with 1 µg/mL CM-DiI membrane lipid stain and 1 µg/µL DAPI. For OMV-RNA co-localisation, samples were stained with 1 µg/mL DiD membrane lipid stain and SYTO RNASelect (0.5 µM). Slides and coverslips were cleaned with 0.02% Hellmanex in an ultrasonic bath for 30 min, rinsed, and dried under a nitrogen stream before being activated in a Tergeo Plus Plasma Cleaner. Images were captured on an Andor Dragonfly 200 (Oxford Instruments, UK) confocal microscope.

### (d) OMV concentration, zeta (ζ) potential, and size measurement

Nanoparticle tracking analysis (NTA) measurements of OMV concentration, ζ potential, and size were performed using a ZetaView x30 Mono (Particle Metrix, Germany). OMV concentrations were recorded as the average of measurements at 11 positions. To determine the number of OMVs per cell, calibration curves were generated by measuring the OD_600_ absorbance of bacterial cultures using spectrophotometry and the corresponding count of cells using flow cytometry. Zeta potential was calculated as the average of three technical replicates each measured at two stationary layers. Determination of OMV size based on electron microscopy imaging was performed using Fiji (ImageJ v1.54p). Briefly, the area of OMVs with a circularity >0.7 was calculated. Each vesicle was converted into an equivalent circle and the diameter calculated. Structures <66 nm were excluded from statistical analysis as they could not reliably be identified as OMVs.

### (e) Statistical analyses

To evaluate co-localisation of OMVs and nucleic acids, the Manders co-localisation coefficient for each image was calculated using the Fiji (ImageJ v1.54p) Coloc2 plug-in. The Costes method was used to determine the likelihood the co-localisation coefficient was a result of chance, by calculating the frequency that a coefficient generated from randomised co-localisation would exceed that of the observed co-localisation [23]. To determine if OMV concentration, ζ potential, or size was correlated with cell density, t-tests of Pearson correlation coefficients were performed. Reported sample concentrations were corrected for culture OD_600_, resuspension volume and dilution factors. Statistical analyses were conducted in RStudio 2024.12.1 Build 563 or with Microsoft Excel 365 v2510.

## 3. Results

### (a) The bee gut microbiota produces OMVs

We examined the nanoscale fraction from liquid cultures of bacterial strains using TEM imaging. All gram-negative members of the core bee gut microbiota (*S. alvi, G. apicola, G. apis, F. perrara*, and *Bartonella apihabitans)* produced near-spherical, membrane-bound structures (Figure 1a, Figure S1). These were consistent, in size and shape, with OMVs described in other bacterial species, including *Pseudomonas aeruginosa, Vibrio cholerae, Klebsiella pneumoniae* and *Xanthomonas campestris* [24–26], as well as with OMVs collected from the hyper-vesiculating *Escherichia coli* strain BW25113 Δ*nlpI* (Figure S1). We did not observe these structures in fractions isolated from the gram-positive bee gut symbiont *L. apis*; gram-positive bacteria can also produce extracellular vesicles, although less frequently and generally through different mechanisms than that of gram-negative bacteria [27]. These structures were also absent in fractions from the gram-negative opportunistic bee gut pathogen *Serratia marcescens* kz11. OMV production has been shown to be temperature-dependent in another, human-derived, *S. marcescens* strain [28]; thus, the cultivation temperature used in our study (designed to mimic *in vivo* conditions) may have been insufficient to trigger OMV production. Additionally, we observed OMV-sized blebs emerging from intact cells of *G. apicola* wkB1 (Figure 1b), with a size and shape similar to the vesicles seen within the extracellular space. The membranes of these emerging blebs were continuous with the outer membranes of the parent cell. Taken together, we conclude that the OMV-like structures observed from the bee gut bacteria are bonafide OMVs.

**Figure 1.**
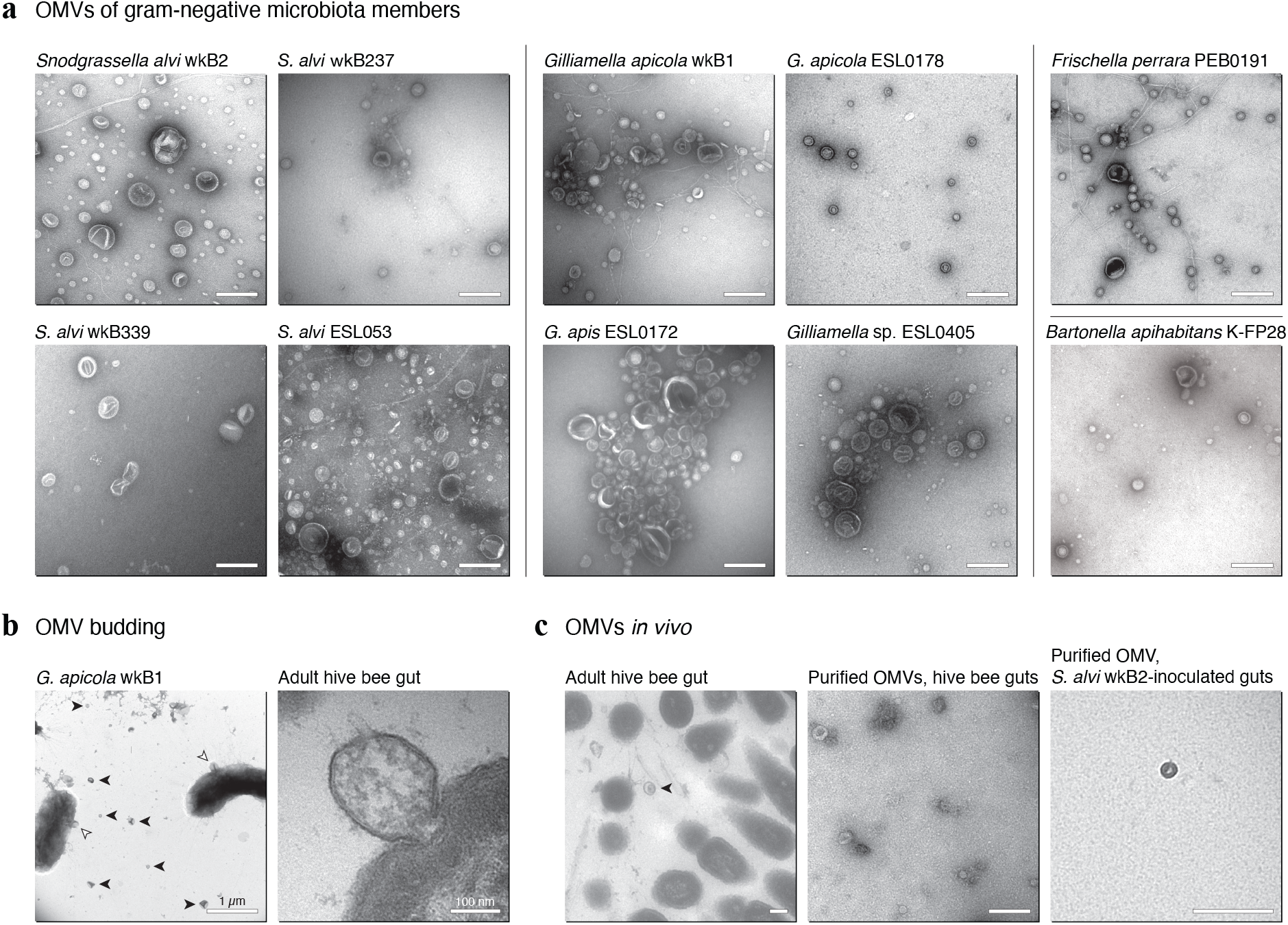
OMVs are produced by bee gut microbiota members *in vitro* and *in vivo*. (a) Representative TEM micrographs of purified OMV fractions from liquid cultures of bee gut bacteria strains showing presence of OMVs from gram-negative members of the normal microbiota. (b) OMVs can be found attached to the bacterial cell surface (unfilled arrows) as well as detached, in extracellular space (solid arrows). (c) Imaging of bee ilea tissue sections show presence of OMVs in wild adult hive bees as well as in lab-reared bees mono-colonised with *S. alvi* wkB2. Scale bars = 200 nm unless otherwise indicated.

### (b) OMVs are produced *in vivo*

Ilea sections from 2-week-old hive bees showed pronounced bacterial colonisation and the presence of putative OMVs identical in appearance to those purified from liquid cultures (Figure 1c). Using TEM tilt tomography on sectioned hive bee ilea, we produced 3D reconstructions showing these vesicles to have a spherical morphology (Electronic Supplementary Material, Video S1). These included vesicles that possess two membranes (Video S2), suggesting that outer-inner membrane vesicles (O-IMVs) [29] may also be produced by the bee gut microbiota. To further verify the existence of OMVs within bee guts, we purified and imaged the liquid contents of pooled ilea from hive bees (pool of 18) or bees mono-colonised with *S. alvi* wkB2 (pool of 22). Vesicles were observed in both the hive bee and *S. alvi*-inoculated ilea (Figure 1c). In contrast, we did not observe any OMV-like structures from two pools of ilea contents (12 and 30 ilea) from non-inoculated bees. NTA measurement showed OMV abundance from hive bee ilea (pool of 18) and *G. apicola* wkB1-inoculated ilea (pool of 21) were 2.8× and 3.3× fold higher, respectively, than from non-inoculated ilea (pool of 30).

### (c) Bee gut bacteria OMVs contain nucleic acid

Purified OMVs were treated with a fluorescent membrane stain in combination with either a fluorescent RNA-specific or DNA-specific stain. To remove nucleic acid not contained within the OMV lumen, samples were treated with RNase or DNase prior to visualisation (Figure S2). Confocal microscopy revealed significant OMV and nucleic acid co-localisation, with Costes *p* < 0.0001 (Figure 2a, 2b). We found the proportion of OMVs containing RNA to be 0.594 (Manders co-localisation, SEM 0.040, n = 12) and DNA to be 0.355 (SEM 0.043, n = 10), with consistently more OMVs containing RNA than DNA (Figure 2c, Figure S2).Extraction of nucleic acid from OMVs collected from 50 mL cultures yielded an average of 146.20 ng (SEM 23.41, n = 32) of RNA and 8.55 ng (SEM 3.26, n = 21) of DNA (Figure 2d).

**Figure 2.**
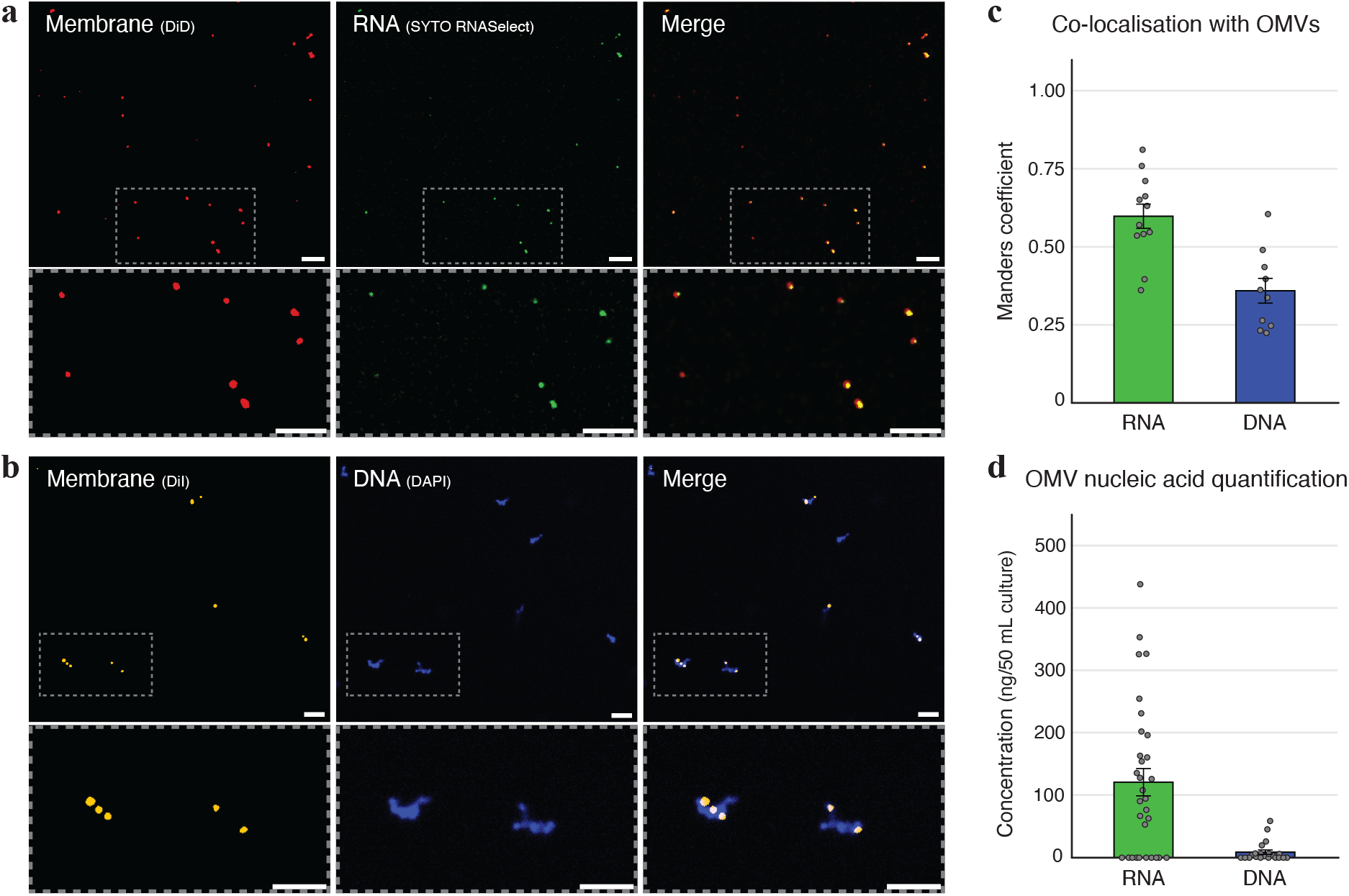
Bee microbiota OMVs contain nucleic acids. Confocal fluorescence imaging of OMVs from *G. apis* ESL0172 shows (a) co-localisation of membrane lipid stain (DiD, red) and RNA stain (SYTO RNASelect, green), and (b) co-localisation of membrane lipid stain (DiI, yellow) and DNA stain (DAPI, blue). Scale bars = 10 μm. (c) Proportion of OMVs with co-localised nucleic acid (Manders correlation coefficient). (d) Quantification of nucleic acids in OMVs from stationary phase cultures. Error bars = SEM.

### (d) OMV characteristics vary by strain and species

NTA was used to estimate OMV concentration, ζ potential, and size. Comparing between strains, we found significant differences in the number of OMVs produced per cell (Figure 3a). *S. alvi* wkB2 produced considerably more OMVs than other strains (mean = 1161.7, SEM = 526.5, median = 260). The maximum mean OMVs per cell for any *Gilliamella* strain was 2.71 (SEM = 1.31, median = 1.40) for ESL0405. This was similar to *B. apihabitans* K-FP28 (mean = 1.72, SEM = 0.30, median = 1.43). OMV concentration was negatively correlated with cell density for all strains (Table S2). Differences were also observed in ζ potential between strains (Figure 3b). *B. apihabitans* K-FP28 and *Gilliamella* sp. ESL0405 had more negative ζ potentials (means < −20 mV), compared to *S. alvi* wkB2, *G. apicola* wkB1, and *F. perrara* PEB0191 (means > −15 mV). For *S. alvi* wkB2 OMVs, size positively correlated with increasing cell density, while OMVs from *Gilliamella* spp. strains and *B. apihabitans* K-FP28 showed weak to strong negative correlations. NTA measures the hydrodynamic diameter of vesicles and can therefore overestimate their size, while conventional TEM involves a degree of dehydration leading to changes in morphology [30]. Therefore, we used cryo-TEM, a technique which preserves hydration, to produce images of *S. alvi* wkB2 and *G. apicola* wkB1 for size analysis (Figure S3). This approach revealed that *S. alvi* and *G. apicola* produces OMVs of different sizes, with median diameters of 76 nm and 95 nm respectively (Figure 3d). A comparison of size by measurement technique is shown in Figure S3.

**Figure 3.**
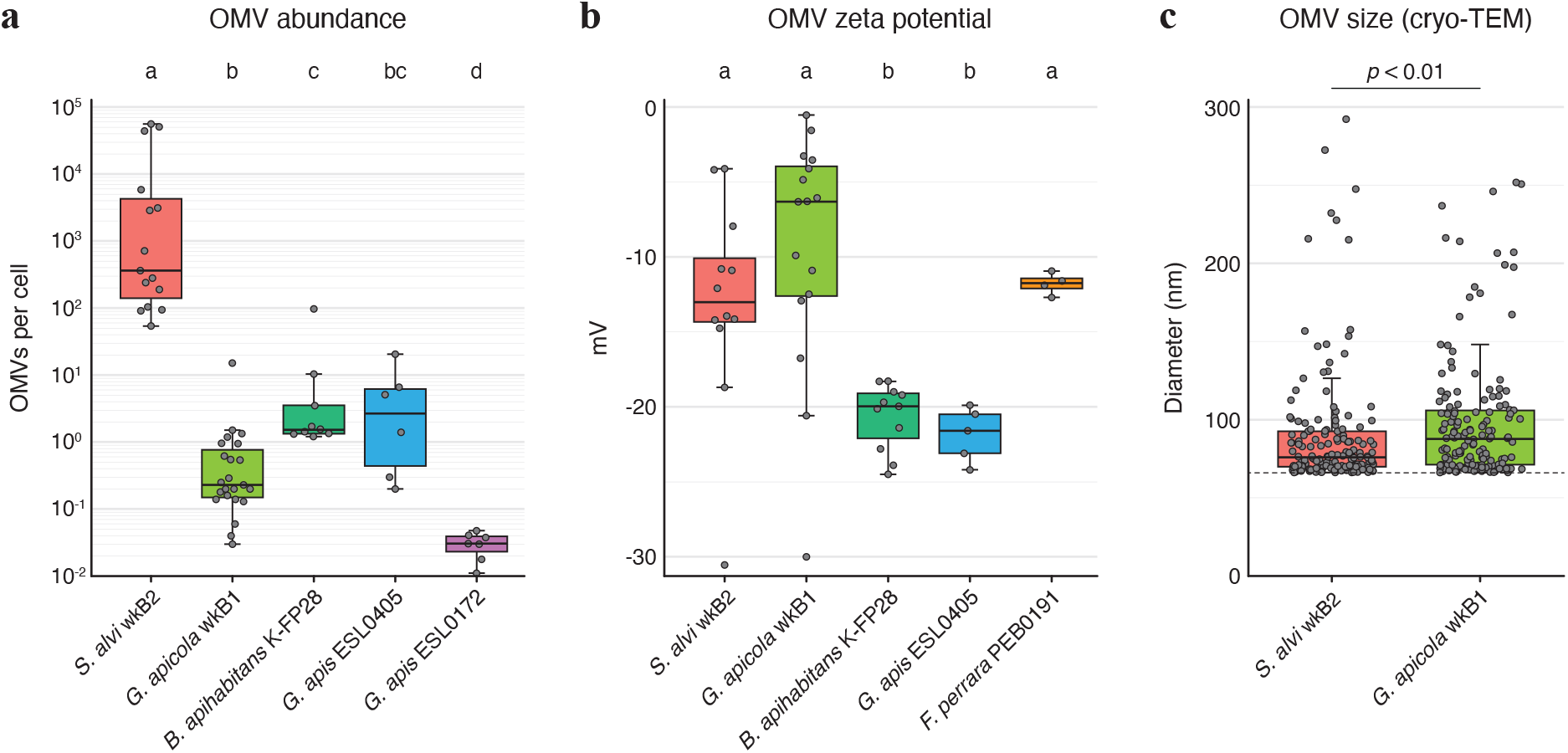
OMV characteristics differ by species. (a) OMV abundance and (b) ζ potential of strains grown in liquid cultures. Significant differences, indicated by letters above boxplots, tested with Welch’s ANOVA and Games-Howell post hoc tests. (c) OMV sizes of *S. alvi* wkB2 and *G. apicola* wkB1, as measured by cryo-TEM. Significance tested with Wilcoxon rank-sum test. Dotted line indicates the size threshold (66 nm) for OMV detection.

## 4. Discussion

OMVs have been widely reported in bacteria, but their role in shaping microbial communities, particularly within symbiotic host-microbe associations, remain poorly understood. We found that the gram-negative core species of the bee gut microbiome produce abundant OMVs. These were generated without induction, in liquid culture and were also observable directly within the ilea in both mono-inoculated bees and in those collected from the wild. Although similar in general appearance, we found substantial differences in concentration and colloidal properties between the tested strains.

*S. alvi* wkB2 produced OMVs at far higher quantities than other strains measured. Several causes of OMV hypervesiculation have been described. These predominantly involve a disruption to the rigidity of the cell envelope; for instance, null mutation phenotypes of cell envelop proteins TolR, DegS, NlpI produce hyper-vesiculating phenotypes in *Salmonella* and *E. coli* [31, 32]. In wildtype strains of *Salmonella*, disruption to cell envelope integrity is often associated to stresses such as antibiotic exposure or acidity [33, 34]. Stress is also induced by increased cell density as nutrients become depleted and waste levels increase [35]. We investigated if this may be the case in our strains by examining if OMV concentration (OMVs/cell) correlated with cell density (Table S2). All five strains we examined, including *S. alvi* wkB2, showed negative correlations with cell density (OD_600_), with three of five being significant (*p* < 0.05). Since the optimal conditions for growth are at the early timepoints of *in vitro* culture, it is unlikely the higher OMV concentrations seen at lower cell densities, or the higher concentrations overall for *S. alvi* wkB2, are stress related.

We observed a positive correlation with cell size and density in *S. alvi* wkB2 and negative correlations for *G. apis* ESL0172 and *Gilliamella* sp. ESL0405. Size variation across growth phases is associated with OMV functional change. For example, in *Helicobacter pylori* it has been shown that OMV size determines the mechanism of entry into host gut epithelia cells [36]. Additionally, the composition of cell envelop proteins in *H. pylori* culture varies with growth phase [37], thereby suggesting a mechanism whereby OMV protein cargo is regulated.

There was wide variation in ζ potential between strains, but this was generally uniform across growth phases (Table S2). Cell surface protein composition is the principal source of ζ potential; therefore, surface charge can change substantially with alterations to cell state and can be variable across growth phases [38]. OMV ζ potential can also vary from that of its parent cell and can have a greater or lesser negative charge [39, 40]. More negative ζ potential contributes to repulsive forces between particles, preventing aggregation and aiding even dispersal through the medium, a process extensively studied in red blood cells [41]. Less negative ζ potential results in lower repulsive force which can lead to increased cell to cell adhesion [42], potentially enabling OMV fusion to target cells and facilitating delivery of cargo.

Luminal RNA and DNA were found in a high proportion of OMVs (Figure 2c), although the amounts of DNA were significantly less than that of RNA. Our preliminary sequencing of OMV-associated DNA from *S. alvi* wkB2 found most of it to be of prophage origin. Co-fractionating with OMVs were other structures of a similar size, including capsids and other phage components, which were noted during TEM imaging (Figure S3b, S3c). Therefore, we cannot discount the possibility that the DNA was derived from phage remnants and not OMV cargo. It is also possible that interactions between OMVs and phages can lead to phage DNA being associated with OMVs. Phage recognition sites have been identified on *Vibrio* and *Salmonella* OMVs, and these have been shown to function in phage resistance via adsorption [9] and DNA ejection [43].

The finding of substantial amounts of OMV associated RNA suggests a possible mechanism for intercellular communication and transcription modulation in the bee gut [44, 45]. OMV-mediated small RNA transport between bacteria and eukaryotic hosts have been described in multiple symbiotic systems, including in the squid light organ [46], plant roots and the human lung [47]. In honey bees, double-stranded RNA from genetically engineered *S. alvi* in the gut is able to translocate into host tissue and trigger an RNAi response [48]. OMVs may be a mechanism for the protection and transport of these molecules, and potentially other factors between gut bacteria and the host.

While we could not directly compare the OMV concentrations between *in vitro* cultures and *in vivo* conditions, qualitatively, there were fewer observable OMVs in ilea tissue sections than in cultures. One contributing factor is that *in vitro* derived OMVs were concentrated from much larger culture volumes than would be present in the gut, resulting in higher apparent vesicle abundance. It is also possible that OMV production or accumulation *in vivo* is moderated by signalling crosstalk between the microbiota and the bee gut epithelium in response to OMV contact, for instance, to facilitate gut colonisation and persistence. OMVs carry microbially derived components (e.g., LPS and peptidoglycan), which can elicit immune responses such as secretion of antimicrobial peptides and reactive oxygen species into the gut lumen [49, 50]. Gut colonisation by symbiotic bacteria is known to be accompanied by modulation of this immune response — a process which can be mediated by OMVs [2, 51]. Such signalling may thereby adapt the local environment to conditions favourable to colonisation and symbiotic host-microbe interactions.

Altogether, our results identify OMVs as pervasive features of the bee gut microbiota, and implicate them as underappreciated mediators of microbial and host interactions within the gut. These findings provide a foundation for future studies of bee bacterial OMVs, with consequences for understanding intercellular communication in beneficial microbiomes and for exploring vesicle-mediated pathways as potential targets for microbiome manipulation.

## Ethics

This work did not require ethical approval from a human subject or animal welfare committee.

## Data accessibility

Electronic supplementary material is available online, including supplementary methods, Figures S1–S3, Tables S1–S3, and Videos S1–S2.

## Declaration of AI use

We have not used AI-assisted technologies in creating this article.

## Authors’ contributions

R.E. and W.K.: conceptualisation, methodology; R.E.: investigation, validation, formal analysis, software, writing—original draft; W.K.: funding acquisition, supervision, Writing – review & editing.

All authors gave final approval for publication and agreed to be held accountable for the work performed therein.

## Acknowledgements

We thank Philipp Engel (Univ. Lausanne), Kasie Raymann (North Carolina State Univ.), and Karina Xavier (Gulbenkian Institute for Molecular Medicine [GIMM]) for providing bacterial strains. We thank the technical support of the Bioimaging Platform of GIMM, funded by PPBI-POCI-01-0145-FEDER-022122, the GIMM Electron Microscopy Facility, the GIMM Flow Cytometry Facility and the Electron Microscopy and X-Rays Facility of the International Iberian Nanotechnology Laboratory.

