## Supplementary Material for "Discovery and characterisation of OMVs produced by the bee gut microbiota"

**Electronic Supplementary Material**

**This file contains:**

Supplementary Methods

Supplementary Figures

Figures S1–S3

Supplementary Tables

Tables S1–S3

Supplementary References

**Additional supplementary files:**

Supplementary Videos

Video S1, S2

### Supplementary Methods

#### *Bacterial culture and OMV collection*

A complete list of strains and culture conditions used in this study can be found in Table S1. For *in vitro* assays, 50 mL liquid cultures were vacuum filtered through 0.2 µm pore membranes (Fisherbrand Cat#FB12566508) to remove cells and large debris, transferred to Beckman Coulter ultracentrifuge tubes (Cat#355642) and spun in a Beckman Coulter Optima LE-80K (SW 28 rotor at 112,400 × *g*) or XE-100 (SW 32 Ti rotor at 120,000 × *g*) ultracentrifuge at 4°C for 5 h. After removing the supernatant, the pellet was resuspended in 1 mL $dH_2O$ and then spun in a 50 kDa molecular weight cut-off filter (Merck Amicon Cat#UFC905008 or NeoBiotech Cat#NB-57-0004-2) at 3,000 × *g* for 6–8 min to remove sub-nano scale material. Retentate (approx. 150 µL per sample) containing the OMVs was collected and stored at 4°C.

For *in vivo* assays, late stage pupae (P6–P7 [1]) were removed from brood chambers, placed into cages, and provided with sucrose water (40% w/v) with or without the addition of *S. alvi* wkB2 at a final concentration of $10^7$–$10^8$ cells/mL. Cages were incubated in the dark at 35°C and 60% relative humidity, for 5–6 days, after which the bees were anaesthetised with $CO_2$ and the ilea excised. Ilea contents were extracted by covering the ilea with 1–2 drops of $dH_2O$ and gently squeezing along its entire length. All extracted contents and ilea were then pooled into 2 mL $dH_2O$ and shaken at 500 rpm at 35°C for 20 min to further dislodge ilea contents with minimal disruption to the ilea tissue. The samples were diluted to 38.5 mL with $dH_2O$ and treated as per the *in vitro* assays.

#### *Electron microscopy sample processing and visualisation*

Gut ileum tissue sections, bacterial cells, and purified OMVs (from bacterial cultures and gut ilea contents) were visualised with transmission electron microscopy (TEM). For visualisation of bacterial cells, 20–50 µL of liquid culture was gently spun down, the media removed and cells resuspended in the same volume of $dH_2O$ before fixation. To produce tissue sections, ilea were fixed for 1 h at room temperature (RT) by immersion in 2% (v/v) formaldehyde and 2.5% (v/v) glutaraldehyde in 0.1 M phosphate buffer, pH 7.4 (PB), followed by 3 × 5 min washes in 0.1 M PB. Sample post-fixation was done with 1% (w/v) osmium tetroxide in 0.2 M PB for 1 h on ice followed by 2 × 5 min washes in 0.1 M PB and 2 × 5 min in $dH_2O$. Samples were stained with 1% (w/v) tannic acid on ice for 20 min, washed 5 × 5 min in $dH_2O$, and immersed in 0.5% (w/v) uranyl acetate (UA) for 1 h at RT in the dark. Samples were then dehydrated sequentially using ethanol dilutions: 30% and 50% EtOH at RT for 10 min; 70% at 4°C overnight; 90% at RT for 10 min and finally 100% at RT for 3 × 10 min. Infiltration with Embed-812 epoxy resin (EPON) was performed sequentially with 3:1 EtOH:EPON; 1:1 EtOH:EPON; 1:3 EtOH:EPON, 90 min each step. These were followed by infiltration of 100% EPON overnight with agitation and a 100% EPON treatment for 30 min. Samples were embedded and hardened overnight at 60°C. Embedded ilea were sectioned using a Leica UC7 Ultramicrotome at a thickness of 70 nm for 2D TEM imaging or

120 nm for tomography and collected onto palladium-copper grids coated with 1% (w/v) formvar in chloroform. Prior to imaging, sections were stained with 1% (w/v) UA and Reynolds' lead citrate for 5 min. 3D tomography imaging was done using a tilt series collected at 23,000× magnification at 1° increments between -55° and 55°. Fiji (ImageJ v1.54p) [2] was used to produce MP4 files. For visualisation of OMVs isolated from liquid cultures and ilea contents, 5 µL of sample was loaded onto glow discharged formvar/carbon 100 mesh copper grids, washed 10× with dH2O and then stained with 2% UA. TEM imaging was performed on a FEI Tecnai G2 Spirit BioTWIN coupled to an Olympus Soft Imaging Solutions Veleta CCD Camera (standard imaging) or an Eagle 4K HS Camera (for tomography) at the Gulbenkian Institute for Molecular Medicine Electron Microscopy Facility.

For cryo-TEM, samples were vitrified with a ThermoFisher Vitrobot Mark IV with the following settings: 100% humidity, blot-force 2, blot-time 2–3 s. Quantifoil® customised R 3.5/1 (3.5 µm holes / 1 µm spacing) 200 mesh copper grids with 2 nm carbon film were used. Grids were made hydrophilic with a 3 s plasma cleaning (Fischione Plasma Cleaner Model 1020 with 3:1 Ar/$O_2$ gas ratio). Cryo-TEM was performed with a Thermo Scientific Glacios microscope at 200 kV, utilising a X-FEG gun, a 12-grid autoloader and a Thermo Scientific Falcon 4i direct electron detector. Imaging used a 100 µm objective aperture, 57k magnification, 2.441 Å pixel size, 22 electron/Å$^2$ dose and a focus range of -3.3 to -2.7 µm. Acquisition was done using Thermo Scientific EPU automatic image acquisition software. Cryo-TEM imaging of vitrified OMVs was performed at the International Iberian Nanotechnology Laboratory.

***Confocal imaging of OMV-associated nucleic acids and co-localisation analysis***

To determine if OMVs contained luminal nucleic acids, we performed fluorescent staining of *G. apis* ESL0172 vesicles for visualisation with confocal microscopy. Prior to staining, vesicle samples were treated with DNA-free™ DNase (Thermo Fisher Scientific Cat#AM1906) and RNase A (Thermo Fisher Scientific Cat#R1253) at a concentration of 100 µg/mL at 37°C for 30 min to remove surface-bound and free-floating nucleic acids. A 700 ng per sample DNA or RNA spike-in, obtained from whole cells, was used to determine digestion effectiveness; this was confirmed by Qubit (Thermo Fisher Scientific) quantification and microscopy. For OMV-DNA co-localisation, OMV samples were stained with 1 µg/mL CM-DiI dye membrane lipid stain (Thermo Fisher Scientific Cat#C7000) and 1 µg/µL DAPI (Thermo Fisher Scientific Cat#62248). For OMV-RNA co-localisation, samples were stained with 1 µg/mL DiD membrane lipid stain (Thermo Fisher Scientific Cat#D7757) and SYTO RNASelect Green (Thermo Fisher Scientific Cat#S32703) at a final concentration of 0.5 µM. Glass slides and coverslips were cleaned with 0.02% Hellmanex (Hellma Cat#9-307-011-4-507) in an ultrasonic bath for 30 min, rinsed with $dH_2O$, and dried under a nitrogen stream before being activated in a Tergeo Plus Plasma Cleaner using oxygen at five standard cubic centimetres per second and a pulse rate of 255 Hz. All images were captured on an Andor Dragonfly 200 (Oxford Instruments, UK) confocal microscope at the Gulbenkian Institute for Molecular Medicine Bioimaging Facility.

Prior to statistical analysis of co-localisation, confocal images were processed as follows. Imaris image (.ims) files were opened in Fiji with default stacking and channel splitting. Brightness/contrast automatic adjustment was applied and each channel converted to 8-bit grayscale. Thresholds were adjusted manually to match signal intensity between channels. Noise and background were reduced after binary conversion using median reduction (radius 3) or “despeckle”. When images showed areas of excessive noise or fluorescent bleaching, the ROI manager was used to define the area of analysis. Images for which signal intensities between channels were not sufficiently equalised were excluded from further analysis. Manders co-localisation coefficient for each image was calculated with Fiji plug-in Coloc2 (Costes regression threshold, 10 randomisations, PSF 3.0). The Costes method [3] was used to determine the likelihood the measured co-localisation coefficient was a result of chance: briefly, we generated a null distribution of Manders coefficients by randomly shuffling pixel values for the nucleic acid channel (1000 permutations) to calculate the frequency that a coefficient generated from randomised co-localisation would exceed that of observed co-localisation.

***OMV nucleic acid extraction and quantification***

Nucleic acid extraction was performed using TRI Reagent (Zymo Cat#R2050-1-200) according to manufacturer’s instructions. Briefly, OMVs collected from stationary phase cultures were lysed by pipetting in TRI Reagent and incubated for 5 min. Chloroform was used for phase separation and recovery was augmented by the addition of 2 µl glycogen (Thermo Fisher Scientific Cat#R0561). Pellets were re-suspended in 10–17 µl nuclease-free water and nucleic acid yields were quantified with a Qubit 2.0 fluorometer using the HS RNA (Thermo Fisher Cat#Q10210) or HS DNA (Thermo Fisher Cat#Q32851) assay kit.

***OMV size, concentration and ζ potential assay and analysis***

Nanoparticle tracking analysis (NTA) measurements of OMV size, concentration and ζ potential was measured using a ZetaView x30 Mono (Particle Metrix, Germany). Measurements were taken at 25°C, a camera frame rate of 7.5, shutter speed of 100 and sensitivity of 70. Zeta ($\zeta$) potential was measured using continuous current with three technical replicates at each of two stationary layers and reported here as an average. Concentrations are reported as the average of a sample measured at 11 positions. All samples, including controls, were diluted with 0.09% PBS to maintain a conductivity between 1000–1500 µS/cm. Acquisition parameters were: minimum brightness 30, minimum area 5, maximum area 1000, trace length 30, ZP /class 1.3, maximum ζ potential 128.1, and no fluorescence filter. Negative controls were generated using uninoculated growth media using the same workflow as for OMV collection. A background signal was observed for tryptic soy broth (TSB). To account for this background, we performed NTA measurements on 22 uninoculated TSB samples.

A limit of blank (LoB) threshold, based on particle concentration, was determined as the mean of TSB background replicates plus 1.645× the standard deviation (one-sided, 95%

confidence). We then set a limit of quantification (LoQ) threshold 10 standard deviations above this value, such that only samples with concentrations equal to, or greater than, the LoQ ($9.2 \times 10^7$ particles/mL) were retained for analysis of any metric. Zeta potential measurements were further filtered to retain only those values representing ≥70% of the total signal for any given sample. This LoQ approach was adapted from standard analytical method validation guidelines [4, 5].

Statistical significances were assessed using either Welch's ANOVA followed by pairwise comparisons using the Games-Howell post hoc test, or, for non-normally distributed datasets, the Wilcoxon rank-sum test. To determine if concentration, size or $\zeta$ potential correlated with cell density, t-tests of Pearson correlation coefficients were performed. All reported sample concentrations have been corrected for culture $OD_{600}$, resuspension volume and dilution factors.

Determination of OMV size based on electron microscopy imaging was performed using Fiji (ImageJ v1.54p). The area of OMVs with a circularity >0.7 was calculated. Each vesicle was converted into an equivalent circle and the diameter calculated. Structures <66 nm were excluded from statistical analysis as they could not reliably be identified as OMVs.

#### *Determination of OMVs per cell*

Bacterial cultures were grown to various densities and $OD_{600}$ was measured using an Absorbance One (Byonoy, Germany) or a DS-11 FX+ (DeNovix, USA) spectrophotometer. The corresponding number of cells were counted using flow cytometry. Briefly, cells were stained with CFSE (Tonbo Biosciences cat#13-0850) according to the manufacturer's instructions, and cell counts were performed on a Cytek Aurora 4-laser flow cytometer with the following settings: FSC 419, SSC 126, SSC-B 137, medium flow rate, 800–1300 events per second and a 30 s stopping time. SpectroFlo software (Cytek, USA) was used for analysis. A calibration equation for determination of cell concentration (cells/mL) at a given $OD_{600}$ was produced using a power-law regression model generated from the flow cytometry derived cell counts. OMVs per cell was calculated as the ratio of OMVs/mL (as measured by NTA) and cells/mL, with the latter estimated by applying the calibration equation to the measured $OD_{600}$ values of the OMV isolation cultures.

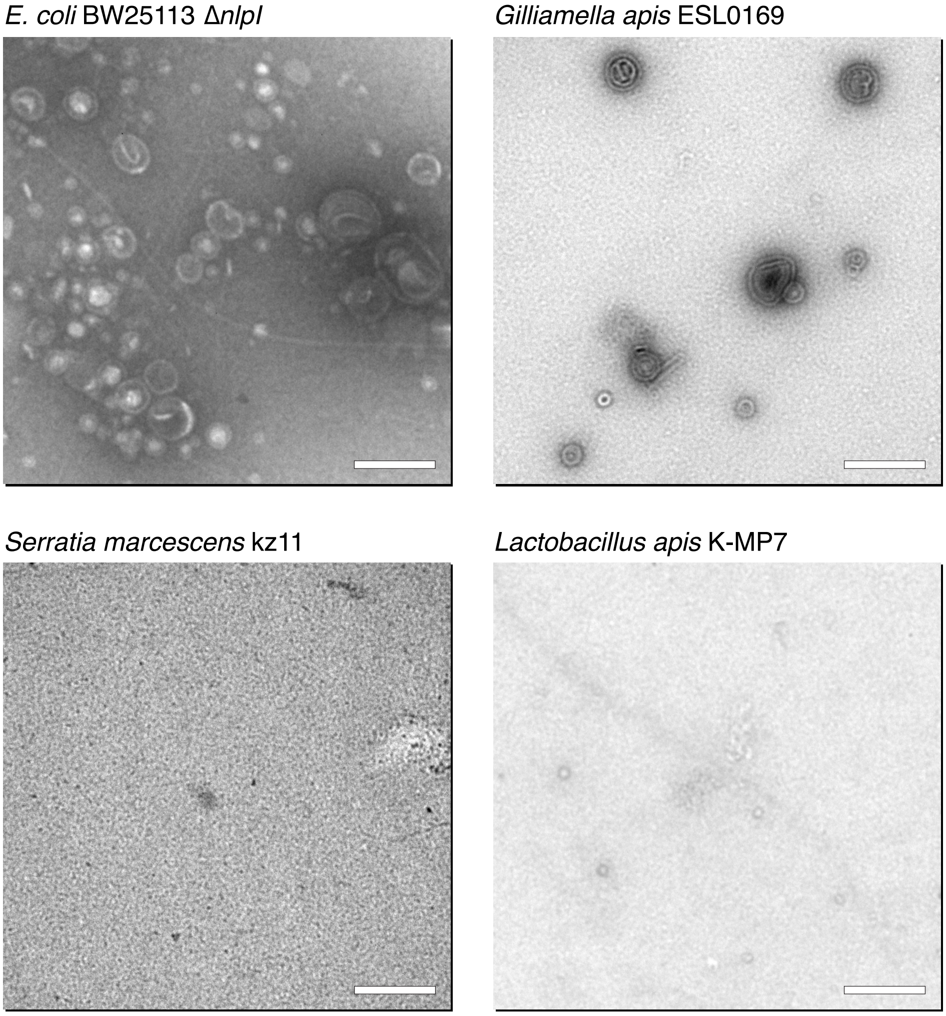


**Figure S1.** Representative TEM images of OMV-purified fractions from a hypervesiculating *E. coli* strain and other bee gut microbiota strains. Scale bars = 200 nm.

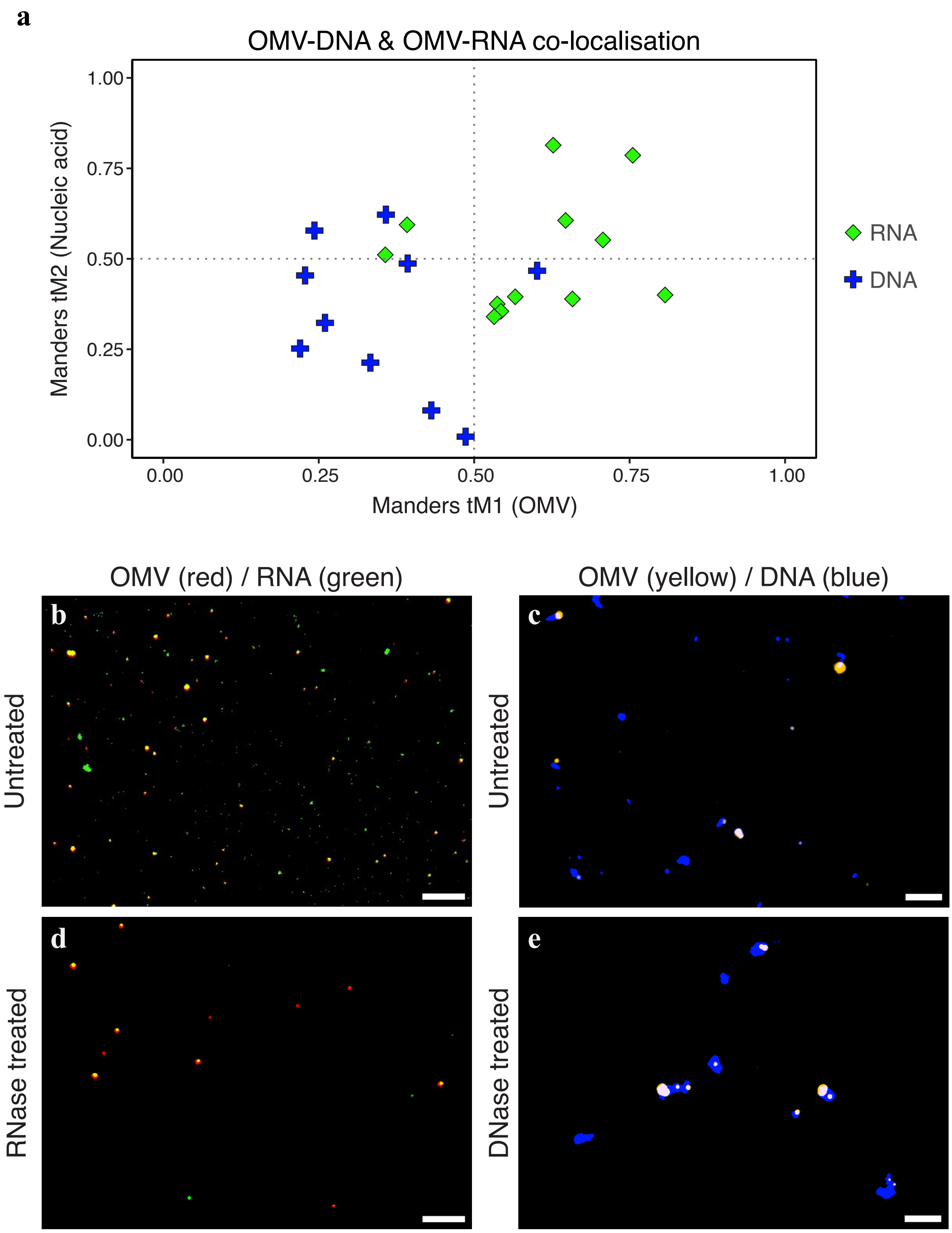


**Figure S2.** Co-localisation of OMVs and nucleic acids. (a) Manders co-localisation coefficient plot (tM1 and tM2) showing imaging replicates with intensities above background threshold. (b–e) Merged confocal images of nuclease treatment controls. OMVs isolated from *G. apis* ESL0172 with the addition of an (b) RNA or (c) DNA spike-in. After treatment with nuclease, a reduction in (d) RNA (green) or (e) DNA (blue) signal was observed. Scale bars = 10 µm.

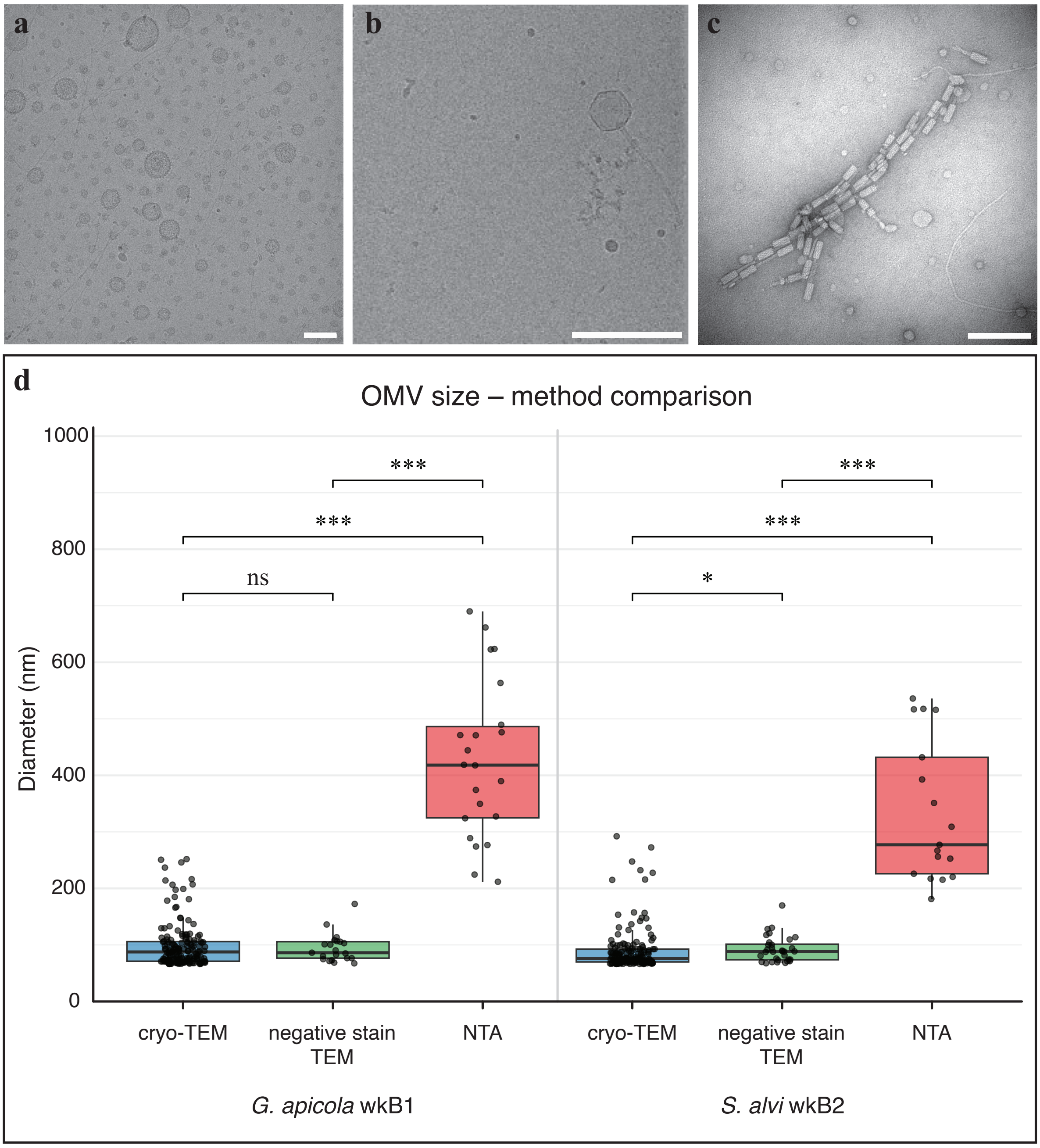


**Figure S3.** Comparison of OMV size measurement methods and images of co-fractionating structures. (a) OMVs from *S. alvi* wkB2 visualised using cryo-TEM. (b) Phage particle from *S. alvi* wkB2 culture observed with cryo-TEM. (c) Possible extracellular contractile injection system structures in *G. apicola* wkB1 culture observed with negative staining TEM. Scale bars = 200 nm. (d) OMV diameter measurements comparing cryo-TEM, negative staining-based TEM and nanoparticle tracking analysis (NTA) methods. Wilcoxon rank-sum test, *p*-values: * <0.05, ** <0.01, *** <0.001, ns = non-significant.

**Table S1**. Strains and culture conditions used in this study. Growth conditions indicate temperature, atmosphere, and shaking rotations per minute (rpm) of liquid cultures.

| Species | Strain | Source | Culture media[1] | Growth conditions |
|---|---|---|---|---|
| *Escherichia coli* | BW25113 Δ*nlpI*::Kan | [6] | LB broth, Miller [7] | 37°C, aerobic, 180 rpm |
| *Serratia marcescens* | kz11 | [8] | Tryptic soy broth (BD Bacto Cat#211825) | 35°C, 5% $CO_2$, 120 rpm |
| *Snodgrassella alvi* | wkB2 | [9] | Tryptic soy broth (BD Bacto Cat#211825) | 35°C, 5% $CO_2$, 120 rpm |
| *Snodgrassella alvi* | wkB237 | [10] | Tryptic soy broth (BD Bacto Cat#211825) | 35°C, 5% $CO_2$, 120 rpm |
| *Snodgrassella alvi* | wkB339 | [11] | Tryptic soy broth (BD Bacto Cat#211825) | 35°C, 5% $CO_2$, 120 rpm |
| *Snodgrassella* sp. | ESL0253 | [10] | Tryptic soy broth (BD Bacto Cat#211825) | 35°C, 5% $CO_2$, 120 rpm |
| *Gilliamella apicola* | wkB1 | [9] | Tryptic soy broth (BD Bacto Cat#211825) | 35°C, 5% $CO_2$, 120 rpm |
| *Gilliamella apicola* | ESL0178 | [12] | Tryptic soy broth (BD Bacto Cat#211825) | 35°C, 5% $CO_2$, 120 rpm |
| *Gilliamella apis* | ESL0169 | [12] | Tryptic soy broth (BD Bacto Cat#211825) | 35°C, 5% $CO_2$, 120 rpm |
| *Gilliamella apis* | ESL0172 | [12] | Tryptic soy broth (BD Bacto Cat#211825) | 35°C, 5% $CO_2$, 120 rpm |
| *Gilliamella* sp. | ESL0405 | [13] | Tryptic soy broth (BD Bacto Cat#211825) | 35°C, 5% $CO_2$, 120 rpm |
| *Lactobacillus apis* | K-MP7 | [14] | MRS broth (VWR Cat#84613.0500) | 35°C, 5% $CO_2$, 120 rpm |
| *Bartonella apihabitans* | K-FP28 | [14] | Heart infusion broth (Merck Cat#05121) + 5% defibrinated sheep blood (Thermo Scientific Oxoid Cat#SR0051C) | 35°C, 5% $CO_2$, 120 rpm |
| *Frischella perrara* | PEB0191 | [15] | Brain heart infusion broth (BD Difco Cat#11708872) | 37°C, anaerobic, 70 rpm |

[1] For growth on solid media, 1.5% w/v agar was added to corresponding broths.

**Table S2.** Correlation of OMV size, concentration, and ζ potential with cell density as measured by nanoparticle tracking analysis. A negative correlation coefficient for size or concentration indicates a decrease in size or concentration with increased cell density. A negative correlation for ζ potential indicates that charge becomes more negative with increased cell density. Concentrations analysed as OMVs per cell. Pearson correlation coefficient with two-tailed *p*-value.

| Species | Strain | n | Comparison | Correlation coefficient | *p*-value |
|---|---|---|---|---|---|
| *Snodgrassella alvi* | wkB2 | 16 | Size | **0.54** | **0.031** |
| *Snodgrassella alvi* | wkB2 | 11 | Concentration | **-0.55** | **0.027** |
| *Snodgrassella alvi* | wkB2 | 11 | ζ potential | 0.02 | 0.957 |
| *Gilliamella apicola* | wkB1 | 22 | Size | -0.26 | 0.236 |
| *Gilliamella apicola* | wkB1 | 22 | Concentration | -0.29 | 0.183 |
| *Gilliamella apicola* | wkB1 | 15 | ζ potential | **-0.51** | **0.046** |
| *Gilliamella apis* | ESL0172 | 7 | Size | **-0.81** | **0.027** |
| *Gilliamella apis* | ESL0172 | 7 | Concentration | **-0.78** | **0.041** |
| *Bartonella apihabitans* | K-FP28 | 10 | Size | -0.45 | 0.197 |
| *Bartonella apihabitans* | K-FP28 | 10 | Concentration | -0.40 | 0.226 |
| *Bartonella apihabitans* | K-FP28 | 7 | ζ potential | -0.08 | 0.870 |
| *Gilliamella* sp. | ESL0405 | 6 | Size | **-0.84** | **0.035** |
| *Gilliamella* sp. | ESL0405 | 6 | Concentration | **-0.92** | **0.009** |
| *Gilliamella* sp. | ESL0405 | 5 | ζ potential | 0.40 | 0.503 |

**Table S3.** Equations used to calculate cell number at various optical densities. Cell numbers were measured by flow cytometry. Values were derived from a power-law regression model (cells/mL = a × $OD_{600}^{b}$). Errors represent the residual standard.

| Species | Strain | n | $R^2$ | Equation ($y = a \times x^b$) | Residual standard error |
|---|---|---|---|---|---|
| *Snodgrassella alvi* | wkB2 | 6 | 0.987 | cells/mL = 2.47e+06 × $OD_{600}^{\wedge 1.231}$ | 8% |
| *Gilliamella apicola* | wkB1 | 9 | 0.954 | cells/mL = 1.2e+09 × $OD_{600}^{\wedge 1.861}$ | 27.7% |
| *Snodgrassella alvi* | wkB339 | 8 | 0.987 | cells/mL = 7.71e+08 × $OD_{600}^{\wedge 3.342}$ | 13.5% |
| *Bartonella apihabitans* | K-FP28 | 8 | 0.889 | cells/mL = 5.15e+08 × $OD_{600}^{\wedge 1.829}$ | 36.3% |
| *Gilliamella* sp. | ESL0405 | 8 | 0.767 | cells/mL = 9.04e+08 × $OD_{600}^{\wedge 1.562}$ | 38.3% |
